# RNA-binding protein Orb2 causes microcephaly and supports centrosome asymmetry in *Drosophila* neural stem cells

**DOI:** 10.1101/2021.11.23.469707

**Authors:** Beverly V. Robinson, Joseph Buehler, Taylor Hailstock, Temitope H. Adebambo, Junnan Fang, Dipen S. Mehta, Dorothy A. Lerit

## Abstract

To maintain a balance of self-renewal versus neurogenesis, neural stem cells (NSCs) undergo asymmetric cell division along an invariant polarity axis instructed by centrosomes. In the NSCs of the third instar *Drosophila* larval brain, the interphase centrosomes are defined by marked asymmetries in protein composition and functional activity as microtubule-organizing centers. Here we show that a conserved RNA-binding protein, Orb2, supports NSC centrosome asymmetry by localizing to the cytoplasm, where it promotes robust apical centrosome maturation and transient basal centrosome inactivation, required for centrosome segregation and spindle morphogenesis. Orb2 is required cell autonomously within NSCs to support centrosome asymmetry and maintenance of the stem cell pool. We suggest Orb2 plays opposing roles in centrosome activation and inactivation at the apical versus basal centrosomes respectively, possibly through the translational regulation of multiple mRNAs. Conversely, loss of *orb2* manifests in microcephaly independent of Orb2 function in NSCs. Bioinformatics uncovers a significant overlap among RNA targets between *Drosophila* Orb2 and human CPEB4, consistent with a conserved role for CPEB proteins in centrosome regulation and neurodevelopment.

## Introduction

*Drosophila* neural stem cells (NSCs) undergo asymmetric cell division (ACD) along an invariant apical-basal polarity axis to segregate cell fate determinants, giving rise to two differentially fated progeny: a self-renewing stem cell and a ganglion mother cell (GMC) destined for neural differentiation (Doe et al., 1991; Knoblich et al., 1995; Kraut et al., 1996; Broadus and Doe, 1997). This balance in NSC self-renewal is critical for neurogenesis, as its deregulation can lead to brain tumors or neurodevelopmental disorders, such as microcephaly (Bond et al., 2002; Cabernard and Doe, 2009). Key to NSC homeostasis are centrosomes, which instruct the division axis and organize the bipolar mitotic spindle required to segregate the pro-stem and pro-differentiation cell fate determinants (Cabernard and Doe, 2009; Januschke and Gonzalez, 2010; Wang et al., 2011).

Centrosomes are microtubule (MT)-organizing centers (MTOC) consisting of a central pair of centrioles surrounded by pericentriolar material (PCM), which recruits the γ-Tubulin (γTub) ring complex required for MT nucleation (Conduit et al., 2015). In most cellular models, centrosomes recruit the robust levels of PCM necessary for MT-nucleating activity just before mitotic onset, a process called centrosome maturation (Gould and Borisy, 1977; Khodjakov and Rieder, 1999). Following mitotic exit, centrosomes shed PCM.

In *Drosophila* NSCs, however, centrosomes are subject to an asymmetric centrosome maturation cycle, wherein the apical (daughter) centrosome recruits PCM and organizes MTs, while the basal (mother) centrosome is transiently inactivated until mitotic onset (Rebollo et al., 2007; Rusan and Peifer, 2007; Conduit and Raff, 2010; Januschke et al., 2011). NSC centrosome asymmetry is implicated in apical-basal spindle pole alignment and centrosome segregation (Januschke and Gonzalez, 2010; Januschke et al., 2013; Lerit and Rusan, 2013; Ramdas Nair et al., 2016). A basic molecular framework required for NSC centrosome asymmetry involves asymmetric localization of Centrobin (Cnb) and Polo kinase (Polo) to the apical centrosome in a mechanism also requiring Wdr62 to promote centrosome maturation (Januschke et al., 2013; Ramdas Nair et al., 2016; Gallaud et al., 2020). Conversely, transient inactivation of the basal centrosome requires Bld10/Cep135, Pericentrin-like protein (PLP), and Polo-like kinase 4 (PLK4/SAK) (Lerit and Rusan, 2013; Singh et al., 2014; Gambarotto et al., 2019). Nevertheless, how centrosome asymmetry is regulated remains incompletely understood.

Intriguingly, *Cnb, Cep135, plp, polo*, and *Wdr62* mRNAs were identified as putative mRNA targets for the RNA-binding protein (RBP) Orb2 through an unbiased crosslinked immunoprecipitation and sequencing (CLIP-seq) study of cultured *Drosophila* S2 cells, suggesting that Orb2 might regulate centrosome asymmetry in NSCs (Stepien et al., 2016). Orb2 is a member of the cytoplasmic polyadenylation element binding (CPEB) protein family orthologous to mammalian CPEB2–4 and implicated in mRNA localization and translational control (Huang et al., 2006; Keleman et al., 2007; Mastushita-Sakai et al., 2010; Hafer et al., 2011). Although prior mutant analysis supports a role for *orb2* in NSC spindle orientation and neuronal specification, whether Orb2 contributes to centrosome regulation is unknown (Hafer et al., 2011).

Here, we identify an NSC-autonomous role for Orb2 in establishing centrosome asymmetry associated with misaligned spindles, supernumerary centrosomes, and NSC loss. We also identify an NSC-independent role for Orb2 in regulating brain size, as *orb2* loss leads to microcephaly. Finally, we examine potential targets of Orb2 and propose a model whereby Orb2-mediated translational regulation contributes to asymmetric centrosome maturation.

## Results and Discussion

### Orb2 localizes diffusely within cycling NSCs

Previous work indicates CPEB proteins localize to centrosomes in *Xenopus* oocytes, embryos, and cultured mammalian cells (Groisman et al., 2000; Eliscovich et al., 2008; Pascual et al., 2020). To investigate whether *Drosophila* Orb2 functions within larval brain NSCs to support centrosome asymmetry, we first examined endogenous Orb2 localization using mouse monoclonal antibodies (Hafer et al., 2011). Although Orb2 localizes to neuronal lineages, subcellular localization of Orb2 is not well defined (Keleman et al., 2007; Hafer et al., 2011). We visualized Orb2 in wild-type (WT) NSCs relative to the centriole marker Asterless (Asl; (Varmark et al., 2007)) and the PCM marker Centrosomin (Cnn; (Megraw et al., 1999; Vaizel-Ohayon and Schejter, 1999)). Orb2 appeared dispersed throughout the cytoplasm with a notable enrichment on the apical centrosome (Fig 1A, *WT*). To ensure specificity of Orb2 signals, we similarly examined Orb2 localization in homozygous *orb2*^Δ36^ (hereafter, *orb2*) NSCs. Unexpectedly, we also noted centrosomal labeling upon staining *orb2* NSCs with Orb2 antibodies (Fig 1A, *orb2*). These spurious signals persisted following pre-absorbing Orb2 antibodies against *orb2* null tissue (Fig. 1B). Consistent with the null nature of this allele, western blotting confirmed Orb2 protein is undetectable within *orb2* larval brain extracts (Fig 1C). Taken together, these data suggest that the apparent centrosomal localization of Orb2 is non-specific.

**Figure 1.**
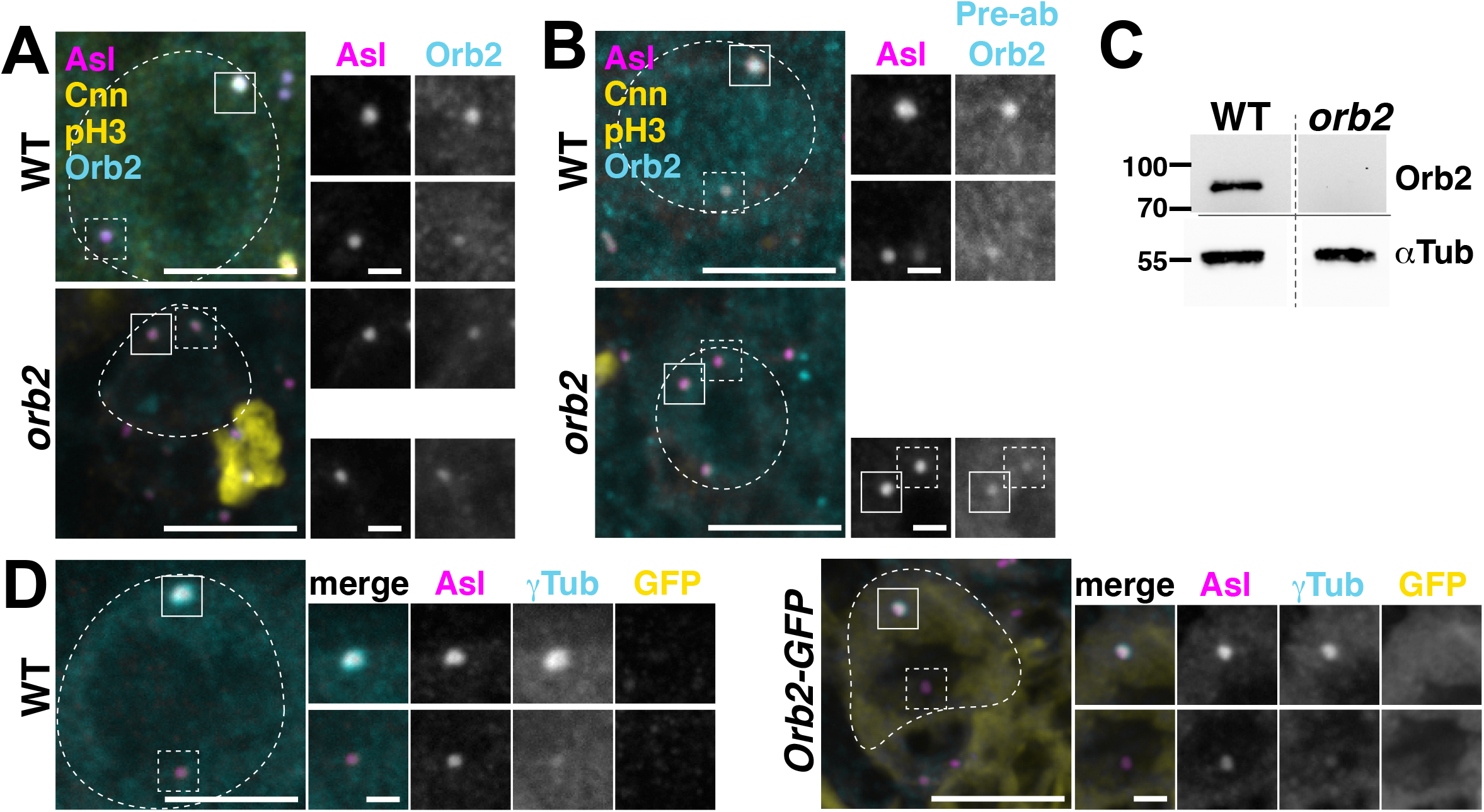
Orb2 localizes diffusely in NSCs. Maximum-intensity projections of (A) WT or *orb2* NSCs (dashed circles) stained for Asl (magenta), Cnn (PCM; yellow), and Orb2 antibodies (cyan). Solid and dashed boxes note apical vs. basal centrosomes; enlarged at right. (B) WT or *orb2* NSCs were stained, as in (A), using pre-absorbed (pre-ab) Orb2 antibodies. (C) Western blot showing Orb2 levels relative to the α-Tubulin loading control in WT vs. orb2 larval brain extracts. The blot was cut along the solid line and lanes were transposed (dashed line). (D) WT or *Orb2-GFP* NSCs stained for Asl (magenta), γTub (PCM; cyan), and GFP antibodies. Bars: 5 μm, 1 μm (inset). Uncropped blots are available to view: https://doi.org/10.6084/m9.figshare.21067519.v1

We next examined the distribution of Orb2 using a transgenic line expressing *Orb2-GFP* under endogenous regulatory elements (Keleman et al., 2007). Staining WT and *Orb2-GFP* brains with anti-GFP antibodies revealed a diffuse distribution of Orb2 throughout the cytoplasm of Orb2-GFP NSCs and GMCs that was absent in WT controls (Fig. 1D). We conclude Orb2 is expressed within cycling NSCs and localizes to the cytoplasm.

### Orb2 disrupts centrosome activity in interphase NSCs

To determine if Orb2 contributes to centrosome asymmetry, we examined γ–Tubulin (γTub) distributions at apical and basal centrosomes in WT vs. *orb2* mutant NSCs during late interphase, when centrosomes are normally asymmetric (Rebollo et al., 2007; Rusan and Peifer, 2007). As anticipated, Asl displayed symmetric distributions among apical and basal centrosomes in both genotypes (mean Asymmetry Index (AI) ± S.D.= 0.0±0.2 for WT and 0.1±0.3 for *orb2*; Fig. 2A–C; n.s. by two-tailed t-test). In comparison, γTub was significantly enriched on the apical centrosome in WT NSCs (Fig. 2A, C). In contrast to WT, *orb2* NSCs showed impaired centrosome asymmetry, evident by increased γTub localization to the basal centrosome and decreased AI values (Fig. 2B, C). γTub AI was significantly reduced by over 25% within *orb2* NSCs, as compared to WT (Fig 2C; *p*<0.01 by two-tailed t-test with Welch’s correction). Consistently, ∼30% of *orb2* NSCs (N= 12/40) had γTub AI values >2 S.D. from the WT mean (Fig 2D; *p*<0.0001 by chi-square test). Measuring the levels of γTub localized at the apical versus basal centrosomes revealed a >2-fold increase in γTub recruitment to the basal centrosomes of symmetrized *orb2* NSCs relative to WT (Fig. 2E; *p*<0.0001 by a two-tailed Mann-Whitney test). In parallel, γTub recruitment to the apical centrosomes trended lower in *orb2* NSCs but was not statistically significant. We conclude Orb2 promotes centrosome asymmetry. Considering Orb2 is dispersed throughout the cytoplasm, Orb2-dependent repression of the basal centrosome during interphase likely occurs from a distance.

**Figure 2.**
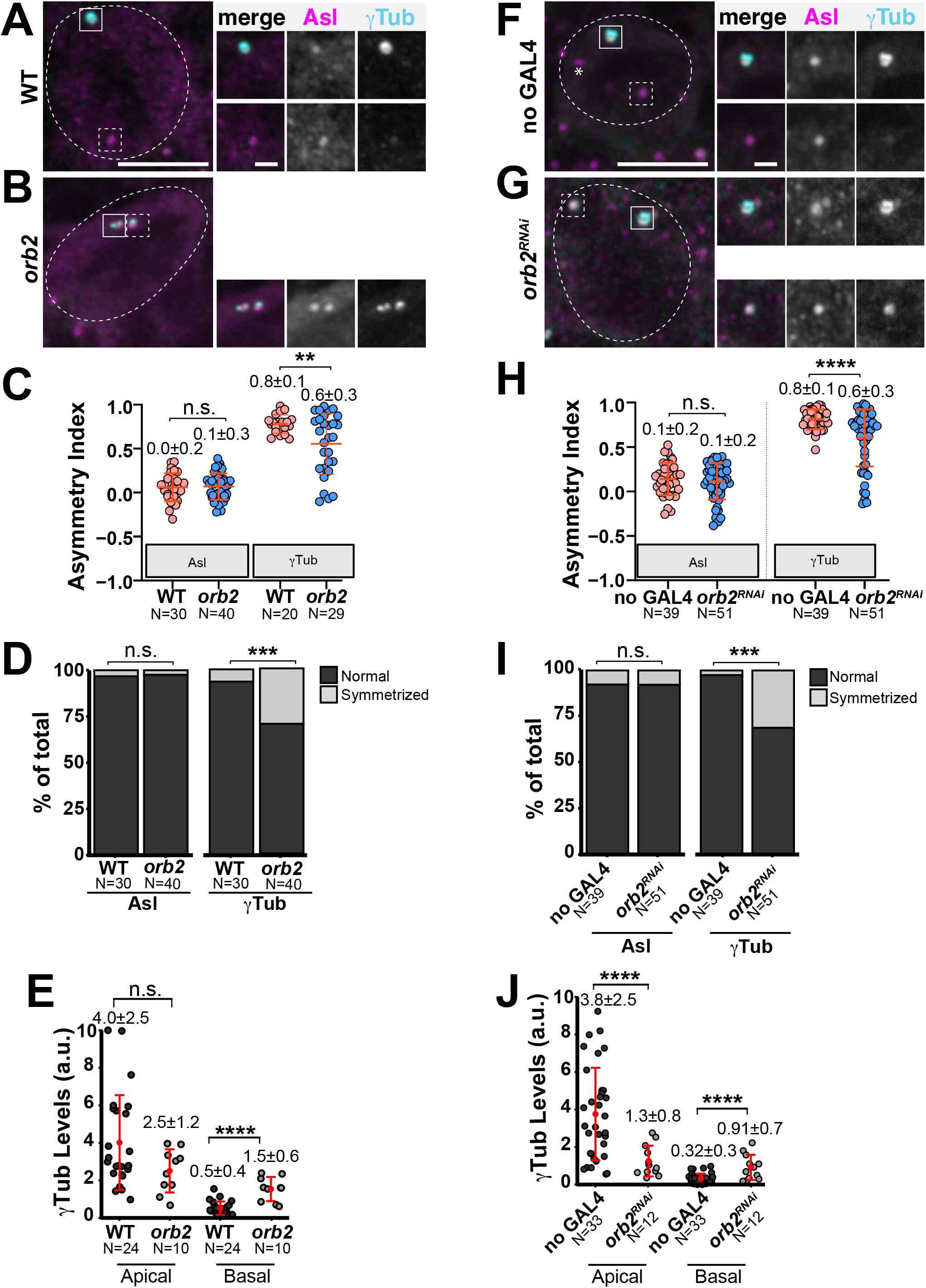
Orb2 contributes to centrosome asymmetry. Maximum intensity projections of interphase NSCs (dashed circles) stained for Asl (centrioles, magenta) and γTub (PCM, cyan). Solid and dashed boxes mark apical vs. basal centrosomes; enlarged at right. (A) WT NSC with γTub enriched at the apical centrosome. (B) *orb2* NSC with γTub at both centrosomes. (C) AI of Asl from N=30 NSCs from n=8 WT brains and N=40 NSCs from n=10 *orb2* brains; AI of and γTub from N=20 NSCs from n=6 WT brains and N=29 NSC from n=8 *orb2* brains. Each dot is a measurement from one cell. (D) Frequency distribution of Asl and γTub AIs in WT vs. *orb2* NSCs. Light grey (symmetrized) values are > 2 S.D. from the control mean. (E) Scatter plot of γTub levels at apical vs. basal centrosomes in N=24 NSCs from n=8 WT brains and N=10 symmetrized NSCs from n=7 *orb2* brains (defined as AI >2 S.D. from WT mean). (F) no *GAL4* control NSCs resemble WT. (G) *orb2*^*RNAi*^ NSC with γTub at both centrosomes. (H) AIs from N=39 NSCs from n=6 no GAL4 brains and N=51 NSC from n=6 *orb2*^*RNAi*^ brains. (I) Frequency distribution of Asl and γTub AIs in control vs. *orb2*^*RNAi*^ brains. (J) Scatter plot of γTub at centrosomes from N=39 NSCs from n=6 control brains and N=16 NSCs from n=6 *worGAL4*>*orb2*^*RNAi*^ brains. Mean ± SD displayed. The experiments were repeated ≥3 independent replicates and significance determined by (C, H) parametric two-tailed t-test, (D, I) chi-square test, (E, J) non-parametric two-tailed Mann-Whitney test: n.s., not significant; *, *p*≤0.05; \*\**p*≤0.01 ***, *p*<0.001; and ****, *p*<0.0001. Bars: 5 μm, 1 μm (insets). Data shown are representative measurements from n>3 independent replicates.

### Orb2-dependent centrosome regulation is cell autonomous

ACD is regulated through intrinsic and extrinsic cellular pathways (Siegrist and Doe, 2006; Doe, 2008). To elucidate if the reduction of centrosome asymmetry observed in *orb2* mutants arose from a requirement for Orb2 within NSCs, we depleted *orb2* specifically in NSCs using an *orb2* dsRNA transgene (*UAS-orb2*^*RNAi*^) driven by the NSC-specific *worniu* (*wor*)*-GAL4* (Albertson et al., 2004). Both the no *GAL4* control and the *wor-GAL4*>*orb2*^*RNAi*^ (hereafter, *orb2*^*RNAi*^) interphase NSCs showed equal distributions of Asl at apical and basal centrosomes (Fig. 2F–H). While controls appeared WT with an enrichment of γTub on the apical centrosome, a subset of *orb2*^*RNAi*^ NSCs (∼30%; N=16/51) recruited significantly more γTub to the basal centrosome and less γTub to the apical centrosome (Fig. 2G–J). These data indicate that Orb2 promotes centrosome asymmetry cell autonomously within NSCs through opposing functions at apical versus basal centrosomes. It is conceivable that Orb2 may modulate PCM recruitment to centrosomes by post-transcriptionally repressing pro-maturation factors (e.g., *Cnb, polo*, or *Wdr62* mRNAs), and/or by promoting the activity of factors required for basal centrosome inactivation (e.g., Cep135 or PLP).

### Loss of *orb2* is associated with supernumerary centrosomes

In *plp* mutants, we previously showed the precocious activation of the basal centrosome is associated with errant centrosome segregation during ACD, resulting in both centrosomes retained within the self-renewing stem cell (Lerit and Rusan, 2013). To assay whether *orb2* loss similarly impairs centrosome segregation, we quantified the frequency of supernumerary centrosomes in WT and *orb2* NSCs. While most WT NSCs (N=25/30) had the expected pair of centrosomes, ∼35% *orb2* mutant NSCs (N=15/40) had extra centrosomes (Fig 3A–C; *p*<0.05 by chi-square test). A similar frequency of supernumerary centrosomes was observed in *orb2*^*RNAi*^ NSCs (∼25%; N=13/51; *p*<0.05 by chi-square test), demonstrating that Orb2 functions within NSC to regulate centrosome number (Fig. 3D–F). Because the two active interphase centrosomes observed upon loss of *orb2* are confined to the apical half of the NSC (Fig. 2B, G), they are likely retained within the stem cell and amplified in the next cell cycle. Alternatively, *orb2* may contribute to centrosome positioning through other mechanisms.

**Figure 3.**
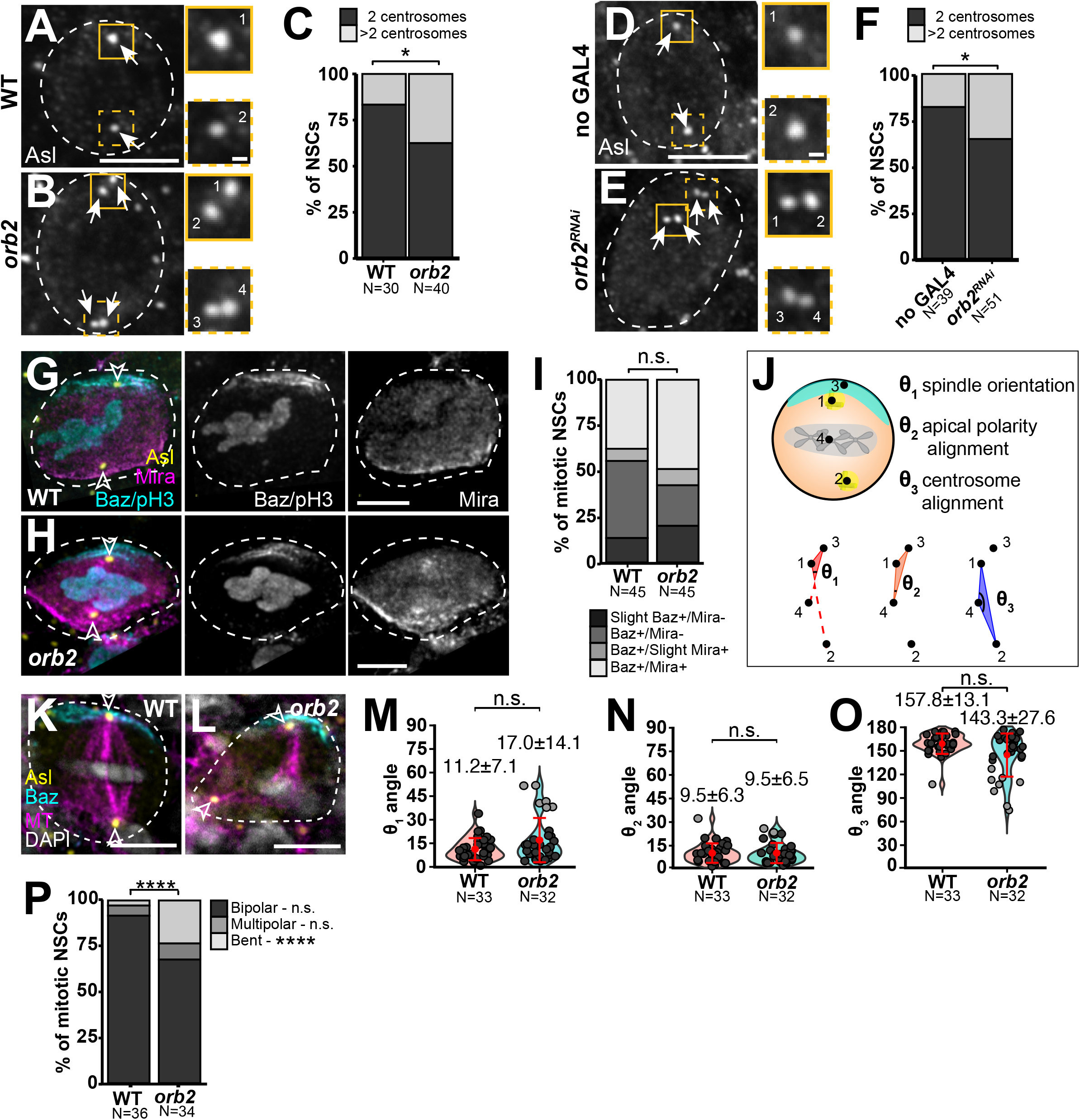
Orb2 is implicated in centrosome segregation and spindle morphogenesis. Maximum intensity projections of interphase NSCs (dashed circles) from (A) WT, (B) *orb2*, (D) no *GAL4* control, and (E) *orb2*^*RNAi*^ stained for Asl (arrows). Solid and dashed boxes mark apical vs. basal centrosomes; enlarged at right. (C and F) Frequency distributions of supernumerary centrosomes in interphase NSCs. (G) WT and (H) *orb2* NSCs stained for Baz (apical cortex, cyan), pH3 (mitotic, cyan), Mira (basal cortex, magenta), and Asl (yellow; arrowheads). (I) Frequency distribution of apical Baz and/or basal Mira crescents in mitotic NSCs. (J) Cartoon depicts points used to measure within mitotic NSCs: θ_1_ spindle orientation, θ_2_ apical polarity alignment, and θ_3_ centrosome alignment (*Methods*). (K) WT and (L) *orb2* NSCs stained for MTs (β-Tub, magenta), Asl (yellow), Baz (cyan), and DAPI (DNA, grey). (M) Plot of θ_1_ spindle orientation. (N) Plot of θ_2_ apical polarity alignment. (O) Plot and (P) frequency distribution of θ_3_ centrosome alignment from N=33 WT and N=32 *orb2* mitotic NSCs from a single experiment across two replicates. Light grey θ are >2 SD from the WT mean. Mean ± SD indicated; significance determined by (M, N, O) unpaired two-tailed t-test or (P) Chi-squared test. n.s., not significant; *, *p*<0.05; and ****, *p*<0.0001. Bars: 5 μm; 1 μm (insets). Data shown are representative measurements from n=3 (A–C) and (K–P) or n=2 (D–F) and (G–I) independent replicates.

### Orb2 is required for mitotic spindle morphogenesis

Given that Orb2 helps regulate centrosome activity and segregation, we next examined its role in spindle orientation. During ACD, the NSC mitotic spindle normally orients along an invariant apical-basal polarity axis entrained by the concerted action of multiple protein complexes (Siegrist and Doe, 2005; Siller et al., 2006; Cabernard and Doe, 2009). Localization of Bazooka (Baz)/Par-3 to the apical cortex in late interphase initiates NSC polarization (Kuchinke et al., 1998; Schober et al., 1999; Wodarz et al., 1999), while localization of the adapter protein Miranda (Mira) to the basal cortex during mitosis represents a late polarization step (Shen et al., 1998; Atwood and Prehoda, 2009). WT and *orb2* mitotic NSCs showed similar distributions of Baz and Mira to the apical and basal cortices, respectively, suggesting that polarization is not significantly disrupted in *orb2* larval NSCs (Fig 3G–I; n.s. by chi-square test).

Prior work identified *αpkc* mRNA as a target for Orb2 translational repression in adult brains and testes (Mastushita-Sakai et al., 2010; Xu et al., 2012; Xu et al., 2014). Further, loss of *orb2* impairs αPKC localization in the testes and embryonic NSCs, the later also showing disrupted spindle orientation (Hafer et al., 2011; Xu et al., 2014). However, our findings indicate larval NSCs polarize correctly in the absence of Orb2, arguing Orb2 may have distinct functions and mRNA targets within the larval brain.

We next used multiple coordinate analysis to define spindle orientation in mitotic WT and *orb2* NSCs stained for β-Tubulin to label MTs, Baz, and Asl (see *Methods*; Fig. 3J–L). First, we measured spindle orientation (θ_1_),, the angle between the apical cortical polarity and spindle alignment axes (Fig. 3J, M). In WT, most spindles aligned within 30° of the polarity axis, yet ∼20% of *orb2* NSC spindles were misoriented >30°, although these differences were not significantly different (Fig 3M).

To ascertain how defects in spindle orientation arise, we next quantified apical polarity alignment (θ_2)_, the angle between the apical centrosome and polarity axes (Fig. 3J, N). Anchoring of the apical centrosome to the apical cortex happens during early interphase and influences polarization in NSCs and other stem cells (Yamashita et al., 2003; Januschke and Gonzalez, 2010; Inaba et al., 2015). Both WT and *orb2* NSCs showed alignment of the apical centrosome to the apical polarity axis, consistent with our observations that apical polarity is unaffected in *orb2* mutants (Fig. 3G-I, N).

Finally, we examined centrosome alignment (θ_3_), the angle between the apical and basal centrosome axes (Fig 3J, O). About 90% of WT mitotic NSCs aligned the basal centrosome ∼180° away from the apical centrosome (Fig 3K, O). In contrast, θ_3_ is reduced in *orb2* NSCs (Fig. 3L, O). While the average reduction in θ_3_ is not statistically significant, a subpopulation of *orb2* NSCs (∼20%; N=7/32 cells) show defective θ_3_ <120° (Fig. 3O, light grey). Consequently, spindle morphogenesis is significantly impaired, resulting in a higher frequency (∼25%; N=9/34 cells; *p*<0.0001 by chi-square test) of bent spindles in *orb2* NSCs relative to WT (Fig 3P). Taken together, these data argue the precocious activation of the basal centrosome in *orb2* NSCs during interphase impairs centrosome migration to the basal cortex, resulting in aberrant spindle alignment and morphology. However, it remains possible that *orb2* may have distinct functions in mitotic spindle orientation.

### Loss of *orb2* results in microcephaly

Defects in centrosome regulation contribute to neurodevelopmental disorders, including microcephaly and intellectual disability (Robinson et al., 2020). Previous work illustrates *orb2* is crucial for learning and memory in the adult *Drosophila* brain (Keleman et al., 2007; Mastushita-Sakai et al., 2010; Kruttner et al., 2012; Hervas et al., 2016). Moreover, reduced brain volumes were noted from serial sectioning adult *orb2* brains (Kruttner et al., 2012). Thus, to assess larval neurodevelopment in *orb2* mutants, we measured the volume of single optic lobes from age-matched third instar larva (Link et al., 2019). Compared to WT brains, *orb2* brains were significantly smaller (Fig 4A, B, and E; *p*<0.0001 by two-tailed t-test). *orb2* brains (N=30/30 brains) had reduced volumes >2-S.D. from WT, consistent with microcephaly.

**Figure 4.**
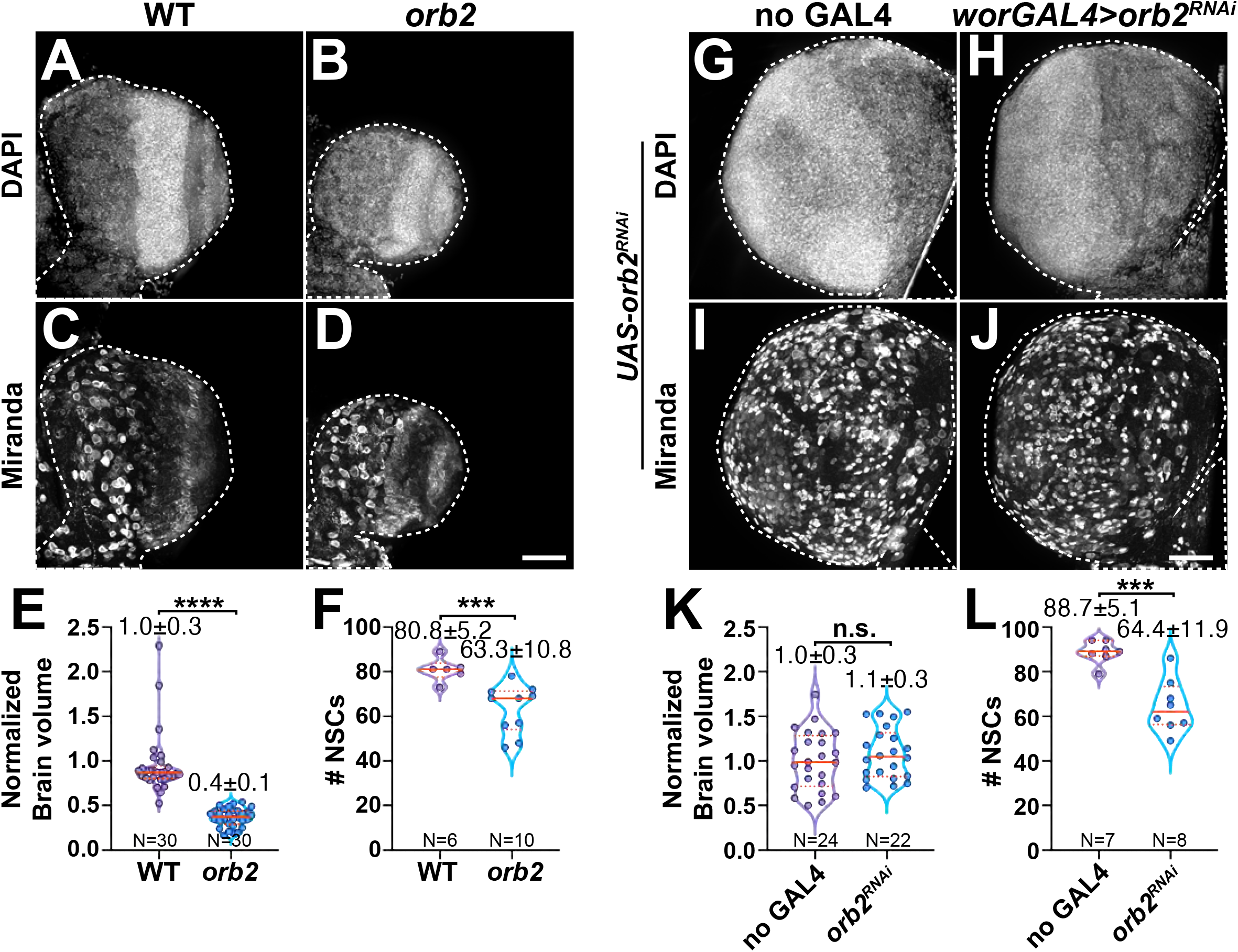
NSC-autonomous and non-autonomous Orb2 activities support neurodevelopment. Projected third instar larval optic lobes (dashed lines) stained for DAPI or Mira. In age-matched brains, (A) WT are larger than (B) *orb2* and (C) WT have more NSCs than (D) *orb2*. (E) Volume quantified from N=30 WT and *orb2* brains. (F) Quantification of NSCs in N=6 WT and 10 *orb2* brains. In age-matched brains, (G) no *GAL4* controls are sized as (H) *orb2*^*RNAi*^ brains. Yet, (I) no *GAL4* controls have more NSCs than (J) *orb2*^*RNAi*^. (K) Volume quantified from N=24 no *GAL4* controls and 22 *orb2*^*RNAi*^ brains and (L) NSC counts quantified from N=7 no *GAL4* controls and 8 *orb2*^*RNAi*^ brains per genotype. Brain volumes were measured from one lobe per brain and normalized to the control mean (Link et al., 2019). Mean ± SD displayed. Significance determined by (E,F, and L) nonparametric two-tailed Mann-Whitney test or (K) unpaired two-tailed t-test; n.s., not significant; ***, *p*<0.001; and ****, *p*<0.001. Bars: 40 μm. Data shown are representative measurements from n=3 independent replicates.

To determine if NSC loss contributes to *orb2*-dependent microcephaly, we counted the number of Mira+ NSCs in control and *orb2* brains. While WT had ∼80 NSCs, *orb2* mutants showed a ∼20% reduction with ∼60 NSCs per lobe (Fig 4C, D, and F; *p*<0.001 by two-tailed t-test). These data indicate Orb2 is required to maintain the NSC pool.

Are the centrosome and spindle defects observed in *orb2* mutants correlated with NSC loss and/or microcephaly? To begin to address this question, we depleted *orb2* specifically in NSCs and compared the volume of age matched *orb2*^*RNAi*^ brains relative to no *GAL4* controls and noted no difference (Fig. 4 G, H, and K), indicating *orb2*-dependent microcephaly is nonautonomous to NSCs – Orb2 is required in other cell types to affect brain size. In contrast, NSC loss is a cell autonomous response to *orb2* depletion. Similar to *orb2* mutants, we detected ∼30% fewer NSC per optic lobe in *orb2*^*RNAi*^ brains relative to no *GAL4* controls (Fig 4I, J, and L; *p*<0.0001 by two-tailed t-test). That the frequency of NSC loss in *orb2* mutants is similar to the incidence of centrosome defects argues these responses are correlated. Moreover, NSC loss is separable from *orb2*-dependent microcephaly. Consistent with the idea that Orb2 supports neurodevelopment in multiple cellular lineages, high levels of Orb2 were previously detected in the cytoplasm of numerous (non-NSC) cells within the larval brain (Hafer et al., 2011).

To elucidate mechanisms underlying NSC loss in *orb2* mutants, we first tested the hypothesis that NSCs are eliminated by cell death. We quantified the coincidence of pro-apoptotic cleaved Caspase 3 (CC3; (Fan and Bergmann, 2010)) in cells marked with the NSC-specific Deadpan (Dpn) antibody (Bier et al., 1992). Similar rates of apoptosis were observed in WT and *orb2* NSCs, indicating NSC loss likely occurs by other mechanisms (Fig S1 A–C; n.s. by t-test). Another mechanism whereby NSCs may be depleted is via premature differentiation (Cabernard and Doe, 2009). Normally, the pro-differentiation marker Prospero (Pros) is confined to the nucleus of the differentiating GMCs and absent from NSCs (Doe et al., 1991; Vaessin et al., 1991), while retention of Pros in the NSC nucleus promotes premature differentiation (Lai and Doe, 2014). We quantified the coincidence of Pros with Dpn and detected no significant difference relative to WT, indicating premature differentiation does not precipitate NSC loss in *orb2* mutants (Fig S1D–F; n.s. by t-test).

Finally, we examined whether *orb2* affects mitotic progression, reasoning impaired NSC self-renewal might contribute to NSC loss. However, the mitotic indices in WT and *orb2* brains were not significantly different (33.6±4.9% per lobe in N=6 WT vs. 36.3±10.1% in N=10 *orb2* brains; Fig S1G–I; n.s. by t-test). These data suggest NSC loss is not due to increased quiescence, although it remains possible that Orb2 may otherwise contribute to cell cycle progression. Alternatively, the altered neuronal specification observed in *orb2* embryos may impinge upon larval neurodevelopment, resulting in fewer NSCs (Hafer et al., 2011).

### Orb2 regulates PLP protein levels in larval brains

Orb2 is an RNA-binding protein known to promote or repress translation of its target mRNAs dependent on its oligomerization status, where monomeric Orb2 functions as a repressor and oligomerized Orb2 functions as a translational activator (Mastushita-Sakai et al., 2010; Xu et al., 2012; Khan et al., 2015). Aspects of the *orb2* null phenotype in NSCs resemble *plp* loss, as both mutants display precocious activation of the basal centrosome in interphase NSCs coincident with supernumerary centrosome and spindle defects (Lerit and Rusan, 2013). Given these similarities, we tested if Orb2 regulates PLP protein expression by comparing PLP levels in WT vs. *orb2* larval brain extracts. Semi-quantitative western blotting uncovered an average 30% reduction in PLP in *orb2* relative to WT, suggesting Orb2 may promote PLP translation (Fig. 5A, B). To assay a requirement for Orb2 in regulating PLP in NSCs, we examined PLP localization to the apical versus basal centrosomes in interphase NSCs. In WT NSCs, PLP was enriched on the inactive basal centrosome, consistent with prior work (Fig. 5C; (Lerit and Rusan, 2013; Singh et al., 2014). Despite a reduction in PLP levels in whole brain extracts, robust localization of PLP to centrosomes was detected in *orb2* NSCs, comparable to WT (Fig. 5C, D; n.s. by two-tailed t-test). Further, levels of PLP localized at apical vs. basal centrosomes were not significantly different from WT (Fig. 5E). Taken together, these data imply Orb2-dependent regulation of PLP may occur outside of NSCs. While the consequences of PLP regulation by Orb2 remain unclear, they are unlikely to cause microcephaly, as *plp* null animals have normal sized brains (Martinez-Campos et al., 2004). We favor the idea that Orb2 likely regulates other mRNA targets to affect centrosome asymmetry and brain volume.

**Figure 5.**
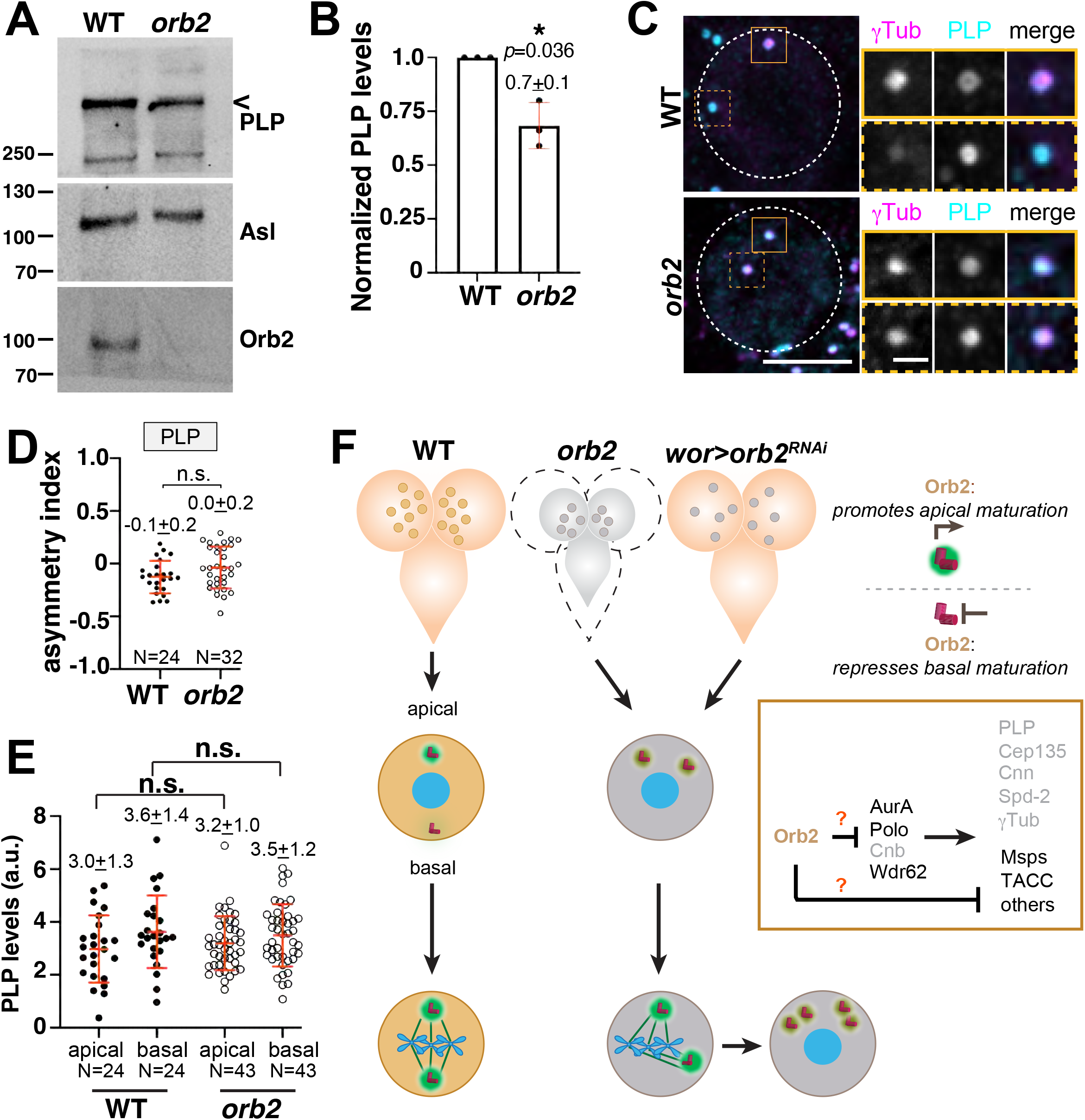
Orb2 regulates PLP protein levels. (A) Immunoblot of PLP, Asl, and Orb2 proteins from WT and *orb2* third instar larval brain extracts. (B) Relative PLP levels were normalized to Asl in WT and *orb2* larval brains from 3 biological replicates. (C) Maximum projected WT and *orb2* NSCs (dashed circles) stained for γTub (magenta) and PLP (cyan). Solid and dashed boxes note apical vs. basal centrosomes; enlarged at right. (D) PLP AI calculated from N=24 WT and 32 *orb2* interphase NSCs. (E) PLP localization to apical and basal centrosomes in N=24 WT vs. N=43 *orb2* interphase NSCs. Data are representative from a single experiment across 2 replicates. Mean ± SD displayed. Significance determined by (B) paired two-tailed t-test, (D and E) unpaired two-tailed t-test; *, *p*<0.05 by ; n.s., not significant. Uncropped, replicated blots are available to view: https://doi.org/10.6084/m9.figshare.17052722 Bars: 5 μm; insets 1 μm. (F) Model depicts the NSC-autonomous function of Orb2 to support interphase centrosome asymmetry and NSC-independent requirement for normal brain volume. Loss of *orb2* activity (grey cells) impairs basal centrosome inactivation and attenuates apical centrosome maturation (PCM, green). We propose Orb2 functions as a translational repressor to support basal centrosome inactivation (*boxed inset;* black text, Orb2 target BC≥3; see Table S1) required for centrosome separation, spindle orientation, and centrosome segregation to the daughter cells. Orb2 may repress the same or distinct mRNA targets to support normal brain volume.

To refine the list of targets that may be involved in centrosome regulation and neurodevelopment, we compared the Orb2 mRNA targets identified in *Drosophila* S2 cells (Stepien et al., 2016) to a recent list of mRNA targets bound by CPEB4 in HeLa cells (Pascual et al., 2020) and identified 1083 overlapping genes (Fig S2A; Table S1). Gene ontology (GO) analysis for cellular components uncovered a significant enrichment of organelle terms (Fig S2B). In contrast, centrosome-related ontologies were not significantly enriched among the overlapping genes (Fig S2C). However, because Orb2 has multiple orthologs, CPEB2-4, other putative Orb2 targets absent from the CPEB4 dataset are omitted from these pairwise analyses. We examined centrosome-related ontologies of Orb2 targets with a biologic complexity (BC) ≥ 3, a metric of reproducibility across biological replicates (Licatalosi et al., 2008). This analysis identified 2150 mRNAs, including 53 mRNAs annotated with centrosome ontologies, which may be subject to Orb2 regulation (Fig. S2D; Table S1). Among these annotated putative Orb2 target mRNAs are *polo, aurora A* (*aurA*), and *Wdr62*, which are known to promote centrosome maturation (Sunkel and Glover, 1988; Giet et al., 2002; Ramdas Nair et al., 2016). One substrate of aurA is transforming acidic coiled-coil protein (tacc), a centrosomal protein that cooperates with mini spindles (msps) to stabilize centrosomal microtubules important for mitotic spindle orientation (Lee et al., 2001). Both *tacc* and *msps* mRNAs are also represented on our annotated list of potential Orb2 targets. Further work is required to determine whether Orb2 RNA-binding activity is required for its functions in regulating centrosome activity, spindle morphogenesis, or brain volume and to subsequently identify relevant RNA targets.

### Proposed Model

Our data indicate Orb2 is required for robust inactivation of the basal centrosome and may also enhance apical centrosome activity in interphase NSCs. We propose the aberrant PCM recruitment in *orb2* NSCs impairs centrosome positioning, resulting in bent mitotic spindles. Moreover, failure to segregate the mispositioned centrosomes among the NSC and GMC produces supernumerary centrosomes in a subset of *orb2* NSCs. Orb2 also functions in other cell types for normal brain size (Fig 5F). While the mechanism of Orb2 function remains unclear, we speculate Orb2 functions remotely in the cytoplasm as a translational regulator to promote apical centrosome maturation (Fig 5F, *top box*) and inactivate the basal centrosome (Fig 5F, *bottom box*). Whether Orb2 functions as a translational repressor or activator to support opposing activities at the apical versus basal centrosomes warrants further study. Based on the apparent molecular weights of bands detected in our immunoblotting experiments, monomeric Orb2B appears to be the predominant isoform within larval brain extracts, given we were unable to detect the smaller Orb2A or higher molecular weight products consistent with oligomerization (Fig 1C and 5A). As monomeric Orb2 functions as a translational repressor (Khan et al., 2015), we predict this is the primary mode of Orb2 activity in larval brains; however, this prediction warrants experimental validation.

Given the uniform distribution of Orb2 within NSCs, one may speculate mRNA targets of Orb2 may themselves be asymmetrically distributed within NSCs. However, patterns of centrosomal RNA localization in NSCs remain to be determined. Alternatively, Orb2 may support opposing functions at the apical versus basal centrosomes by repressing spatially segregated factors. Other models are also feasible, particularly if Orb2 limits levels of a kinase with multiple substrates, any of which could be themselves asymmetrically localized.

Microcephaly is heterogenous; however, a significant number of genes associated with heritable microcephaly are also associated with centrosome biogenesis and regulation (Thornton and Woods, 2009; Megraw et al., 2011; Robinson et al., 2020). Our study implicates Orb2 at the intersection of these pathways.

## Materials and Methods

### Fly Stocks

The following strains and transgenic lines were used: *y*^*1*^*w*^*1118*^ (Bloomington *Drosophila* Stock Center (BDSC) #1495) was used as the WT control unless otherwise noted. *orb2* brains were isolated from the homozygous null allele, *orb2*^*Δ36*^ (P. Schedl, Princeton University (Xu et al., 2012)). NSC-specific depletion of *orb2* by *orb2*^*RNAi*^ (*P{TRiP*.*HMJ22715}*^*attP40*^; BDSC #60424) was driven by *(P{wor*.*GAL4*.*A}*^*2*^; BDSC #56555). All strains were maintained on Bloomington formula cornmeal-agar media (Lab-Express, Inc.; Ann Arbor, MI) at 25°C in a light and temperature-controlled incubator.

### Immunofluorescence

Crawling third instar larval brains were used for dissections and prepared for immunofluorescence as previously described (Lerit et al., 2014). Brains were dissected in Schneider’s *Drosophila* Medium (ThermoFisher Scientific, #21720024), fixed in 9% paraformaldehyde for 15 min, blocked in PBT buffer (Phosphate Buffered Saline (PBS) supplemented with 1% BSA and 0.1% Tween-20) for one hour at room temperature prior to overnight incubation in primary antibodies in PBT with nutation at 4°C. Samples were further blocked with modified PBT (2% BSA, 0.1% Tween-20, and 4% normal goat serum in PBS) for one hour before incubation for two hours at room temperature with secondary antibody and DAPI. Brains were oriented and mounted in Aqua-Poly/Mount (Polysciences, Inc) prior to imaging.

The following primary antibodies were used: guinea pig anti-Asl (1:4000, gift from G. Rogers, University of Arizona), mouse anti-GTU88 (γTub; 1:250–350, Sigma T6557), rabbit anti-Cnn (1:4000 gift from Tim Megraw, Florida State University), guinea pig anti-PLP (1:4000, gift from Nasser Rusan, NIH), rabbit anti-phospho-Histone H3 Ser10 (pH3; 1:2000, Sigma-Millipore, 05-570), mouse anti-β-Tub (clone E7, 1:250; Developmental Studies Hybridoma Bank (DSHB)), mouse anti-Orb2 (undiluted 1:1 mix of clones 2D11 and 4G8 (DSHB); (Xu et al., 2012)), rat anti-Mira (1:500, Abcam, ab197788), rabbit anti-Baz (1:2000, gift from A. Harris, University of Toronto), rat anti-Dpn (1:500, Abcam, ab195173), rabbit anti-cleaved Caspase 3 (CC3, 1:75; Cell Signaling Technology, 9661s), mouse anti-Pros (clone MR1A, 1:500; DSHB), and rabbit anti-GFP (1:700; Proteintech, 50430-2-AP). Secondary antibodies: Alexa Fluor 488, 568, and 647 (1:500, Molecular Probes). DAPI (ThermoFisher Scientific) was used at 10 ng/mL.

### Microscopy

Images were acquired on a Nikon Ti-E inverted microscope fitted with a Yokogawa CSU-X1 spinning disk head (Yokogawa Corp. of America), Orca Flash 4.0 v2 CMOS camera (Hamamatsu Corp.), Perfect Focus system (Nikon), and a Nikon LU-N4 solid-state laser launch (15 mW; 405, 488, 561, and 647 nm) using the following Nikon objectives: 100x 1.49-NA Apo Total Internal Reflection Fluorescence oil immersion, 40x 1.3-NA Plan Fluor oil immersion, and 2x× 0.75-NA Plan Apo. Images were acquired at 25°C through Nikon Elements AR software on a 64-bit HP Z440 workstation (Hewlett-Packard).

### Image analysis

Images were assembled using Fiji (National Institutes of Health; (Schindelin et al., 2012)), Adobe Photoshop, and Adobe Illustrator software to separate or merge channels, crop regions of interest, generate maximum-intensity projections, and adjust brightness and contrast.

#### Centrosome asymmetry

Interphase NSCs were identified by the absence of pH3, round nuclear morphology, and presence of duplicated centrosomes in large (≥10 μm) cells. To blind the experimenter to genotype, maximum projected images were generated, randomized, and used to measure background-subtracted integrated densities from regions of interest (ROIs) centered at the apical or basal centrosomes. Apical centrosome integrated density (*A*) and basal centrosome integrated density (*B*) were used to calculate the asymmetry index (AI), (*A-B*)/(*A+B*) (Lerit and Rusan, 2013). Because the difference between apical and basal integrated densities is divided by the sum, AI is a unitless measure wherein values close to +1 or -1 indicate apical versus basal enrichment, respectively, and 0 indicates an equal distribution. The mean ± S.D. were calculated per genotype.

#### Polarity, NSC number, and mitotic index

Mitotic NSCs were identified by the presence of pH3 in large, Mira+ cells. To score polarity, maximum projected images were anonymized to blind the experimenter to genotype, and each NSC was scored for the absence or presence of Baz or Mira crescents at the apical or basal cortices, respectively. All Mira+ NSCs were counted to quantify the total number of NSCs per optic lobe. Mitotic index is defined as the number of Mira+, pH3+ NSCs per total Mira+ NSCs.

#### Spindle morphology

Z-stack images of mitotic NSCs labeled with anti-β-Tub to label the mitotic spindle, Asl to mark the centrioles, and Baz to label the apical polarity axis were randomized to blind the experimenter to genotype. The point tool in Fiji was used to record the x,y, and z coordinates of four ROIs per cell: 1) center of the apical centrosome, 2) center of the basal centrosome, 3) center of the Baz apical crescent, and 4) center of the DAPI+ condensed chromosomes. These points were used to calculate four different vectors, and vector analysis was used to calculate the angles between specified vectors. The following angles (*θ*) were calculated: *θ*_1_= *spindle orientation*, the angle between the vectors defined by points 1 and 2 (division axis) relative to points 3 and 4 (polarity axis); *θ*_2_= *apical polarity alignment* is the angle between the vectors defined by points 3 and 4 (apical polarity axis) relative to points 1 and 4 (apical centrosome axis); and *θ*_3_= *centrosome alignment* is the angle between the vectors defined by points 1 and 4 (apical centrosome axis) relative to points 2 and 4 (basal centrosome axis). Angles that fell outside ± 2 S.D. from the control mean were defined as defective, as indicated with light grey dots on the corresponding scatter plots.

#### Age-matched brain volume

To synchronously stage third instar larva, *Drosophila* adults were allowed to seed grape juice agar plates supplemented with yeast paste for 48 hrs at 25°C. Plates were changed every 24 hrs, after which flies were allowed 30 minutes to clear previously fertilized eggs. Embryos were collected for four hrs on a fresh plate and incubated at 25°C until first instar larva hatched. First instar larva were transferred into vials containing *Drosophila* culturing medium supplemented with yeast paste dyed with bromophenol blue, then incubated at 25°C until crawling third instar larva emerged. Age-matched larva that both displayed foraging behavior and partial gut clearance of the blue yeast paste were then selected for dissection (Truman and Bate, 1988).

Volume measurements were obtained in Imaris 9.9 software (Oxford Instruments) using the 3D surface tool to define the optic lobe region of interest, visualized with DAPI, and statistics function to calculate surface volume (Link et al., 2019). The experimenter was blinded to genotype prior to analysis.

#### NSC differentiation and death rate

NSCs were identified by the presence of Dpn. To score for premature differentiation, maximum projected images of Dpn and Pros stained brains were randomized to blind the experimenter to genotype, and an ROI of the central brain region was used to calculate the Pearson’s Correlation Coefficient on background-subtracted and automatic threshold–masked images using the Coloc 2 plugin for Fiji (Schindelin et al., 2012). To score cell death, a similar analysis was run on Dpn and CC3 stained brains. To assess specificity of overlapping signals for both experiments, the red channel (Pros or CC3) was rotated 90° clockwise, and colocalization was remeasured.

### Immunoblotting

Larval brain extracts were prepared from 10-20 crawling third instar larva dissected in Schneider’s medium, removed of imaginal discs, transferred to fresh media, then rinsed once in cold PBST (PBS plus 0.1% Tween-20). Samples were homogenized on ice in 30 μL of fresh PBST using a cordless motor and plastic disposable homogenizer, supplemented with 20 μL 5x SDS loading buffer, boiled for 10 min at 95 °C, then stored at -20 °C or resolved on a commercial 7.5% polyacrylamide gel (Bio-Rad, #4568023). Proteins were transferred to a 0.2 μm nitrocellulose membrane (GE Healthcare) by wet transfer in a buffer containing 25 mM Tris-HCl, pH 7.6, 192 mM glycine, 10% methanol, and 0.02% SDS at 4 °C. Membranes were blocked in 5% dry milk in TBST (Tris-buffered saline, 0.05% Tween-20), washed with TBST, and incubated overnight at 4 °C with primary antibodies. After washing with TBST, membranes were incubated for 1.5 hr in secondary antibodies diluted 1:5000 in TBST. Bands were visualized with Clarity ECL substrate (Bio-Rad, 1705061) on a Bio-Rad ChemiDoc imaging system.

The following primary antibodies were used: rabbit anti-PLP (1:4000, gift from Nasser Rusan, NIH), guinea pig anti-Asl (1:10,000, gift from Greg Rogers, University of Arizona), mouse anti-Orb2 (1:25 dilution each of 2D11 and 4G8; DSHB, Paul Schedl, Princeton University (Hafer et al., 2011)), and mouse anti-α-Tubulin (1:10,000, Sigma-Aldrich DM1α). Secondary antibodies: goat anti-mouse HRP (1:5000, ThermoFisher #31430), goat anti-rabbit HRP (1:5000, ThermoFisher #31460), and goat anti-guinea pig HRP (1:5000, ThermoFisher #A18769). Densitometry was measured in Adobe Photoshop and protein levels are normalized to the Asl loading control. Full-size, replicated blots are available to view on FigShare: https://doi.org/10.6084/m9.figshare.17052722.

### Bioinformatics

Gene names were converted to FlyBase identifiers using the Flybase.org tool ‘Query by symbols’ (Larkin et al., 2021). Overlapping genes were identified by the ‘COUNT IF’ function in Excel and Venn diagrams were plotted in R-Studio. GO cellular component analysis was done using the Panther statistical overrepresentation test (http://www.pantherdb.org/), and Fisher’s exact test was used to generate an adjusted p-value, i.e., false discovery rate (FDR) (Mi et al., 2021).

### Statistical analysis

Data were plotted and statistical analysis performed using Microsoft Excel, GraphPad Prism, and RStudio software. Normality distributions were determined using a D’Agnostino-Pearson or Shapiro-Wilk normality test. Goodness-of-Fit chi-squared analysis were performed using control distributions as the expected distributions. Otherwise, data were analyzed using a parametric unpaired two-tailed t-test. In cases where the S.D. were different, we also applied a Welch’s correction. For non-normally distributed data, we used a nonparametric two-tailed Mann-Whitney test. Data are plotted as mean ± S.D. and are representative results from at 2 or more independent experiments, as described in the legends.

## Supplemental materials

**Fig. S1 NSC differentiation and survival in WT vs. *orb2* mutants**. Maximum intensity projections of (A) WT and (B) *orb2* brains stained with Dpn (green; NSC nuclei) and CC3 (red; pro-apototic. Insets (boxes) are enlarged to the right to highlight NSCs (dashed ovals). (C) Quantification of the Pearson’s correlation coefficient between Dpn and CC3 from N=10 WT and 15 *orb2* optic lobes pooled from two replicates. The CC3 channel was rotated clockwise (90°) to test for specificity of overlapping signals. (D) Maximum intensity projections of WT and (E) *orb2* brains stained with Dpn (green) and Pros (red; differentiated nuclei). (F) Quantification of the Pearson’s correlation coefficient between Dpn and Pros from N=16 WT and 18 *orb2* optic lobes pooled from two replicates. The Pros channel was rotated clockwise (90°) to test for specificity of overlapping signals. In (D) and (F), each data point is from one ROI per optic lobe. (G) Projected images of WT and (H) *orb2* NSCs (dashed ovals) stained for Mira (green; NSCs) and the mitotic marker pH3 (red). (I) Quantification of mitotic index from N=6 WT and 10 *orb2* brains. n.s., not significant by Student’s t-test. Bars: (A–E) 20 μm and 8 μm (insets); (G and H) 10 μm.

**Fig. S2 Ontological analysis of Orb2 and human CPEB4**. A) Venn diagram showing relative transcript pool sizes and overlap (green) detected from the Stepien et al. Orb2 CLIP (light blue) and Pascual et al. CPEB4 (orange) datasets (Pascual et al., 2020). B) Top 5 gene ontologies overrepresented in the overlapping transcripts by lowest raw *p* value. C) Overrepresentation tests for gene ontologies implicated in the centrosome were not significant. D) Overrepresentation test of genes in the Orb2 CLIPseq with BC *≥*3 yields significant centrosome ontologies. -log *p*-values are plotted with a significance cut-off of *p*=0.05, noted by the dashed line and asterisk. Genes are listed in Table S1.

**Table S1. Orb2 and CPEB4 common RNA targets**. Sheet 1 lists FlyBase IDs for all transcripts detected in (Pascual et al., 2020). Sheet 2 lists overlapping genes; column E is sortable to list genes with centrosome ontologies. Sheet 3 lists GO terms overrepresented in the overlapping genes, column J is sortable to list genes with centrosome ontologies. Sheet 4 lists Orb2 targets from (Stepien et al., 2016) with BC ≥ 3; column F is sortable to list genes with centrosome ontologies. Sheet 5 shows an overrepresentation test for centrosome ontologies from the Orb2 BC ≥ 3 gene list. Sheet 6 lists each gene included in the indicated centrosome ontologies.

## Acknowledgements

We thank Tony Harris, Timothy Megraw, Nasser Rusan, Greg Rogers, and Paul Schedl for gifts of reagents. We are grateful to members of the Lerit lab, Victor Faundez, and Nasser Rusan for constructive feedback on the manuscript.

This work was supported by NIH grants T32GM008490 (BVR), T32 GM135060-03 (JB and TH), K99GM143517 (JF), and R01GM138544 (DAL).

## Author contributions

Beverly V. Robinson– data curation, formal analysis, investigation, methodology, visualization, and writing–original draft.

Joseph D. Buehler– investigation, formal analysis, software, visualization, and writing–review & editing.

Taylor S. Hailstock – investigation, formal analysis, visualization, writing – original draft, and writing–review & editing.

Temitope H. Adebambo – investigation, formal analysis, and writing–review & editing. Junnan Fang– formal analysis, funding acquisition, investigation, and methodology.

Dipen S. Mehta– investigation

Dorothy A. Lerit– conceptualization, formal analysis, funding acquisition, investigation, methodology, project administration, supervision, visualization, writing–original draft, and writing–review & editing.

## Abbreviations used

ACD: asymmetric cell division
AI: asymmetry index
Asl: Asterless
AurA: aurora A kinase
Baz: Bazooka/PAR-3
BC: biologic complexity
Cep135: Centrosomal protein 135kDa
Cnb: Centrobin
Cnn: Centrosomin
γTub: γ-Tubulin
GMC: ganglion mother cell
Mira: Miranda
Msps: mini spindles
MT: microtubule
MTOC: microtubule-organizing center
NSC: neural stem cell
PCM: pericentriolar material
PLP: Pericentrin-like protein
Polo: Polo kinase
Tacc: transforming acidic coiled coil
Wdr62: WD repeat domain 62
Wor: worniu WT, wild-type

## References

Albertson, R., C. Chabu, A. Sheehan, and C.Q. Doe. 2004. Scribble protein domain mapping reveals a multistep localization mechanism and domains necessary for establishing cortical polarity. J Cell Sci. 117:6061–6070.

Atwood, S.X., and K.E. Prehoda. 2009. aPKC phosphorylates Miranda to polarize fate determinants during neuroblast asymmetric cell division. Curr Biol. 19:723–729.

Bier, E., H. Vaessin, S. Younger-Shepherd, L.Y. Jan, and Y.N. Jan. 1992. deadpan, an essential pan-neural gene in Drosophila, encodes a helix-loop-helix protein similar to the hairy gene product. Gene Dev. 6:2137–2151.

Bond, J., E. Roberts, G.H. Mochida, D.J. Hampshire, S. Scott, J.M. Askham, K. Springell, M. Mahadevan, Y.J. Crow, A.F. Markham, C.A. Walsh, and C.G. Woods. 2002. ASPM is a major determinant of cerebral cortical size. Nature genetics. 32:316–320.

Broadus, J., and C.Q. Doe. 1997. Extrinsic cues, intrinsic cues and microfilaments regulate asymmetric protein localization in Drosophila neuroblasts. Curr. Biol. 7:827–835.

Cabernard, C., and C.Q. Doe. 2009. Apical/basal spindle orientation is required for neuroblast homeostasis and neuronal differentiation in Drosophila. Dev. Cell. 17:134–141.

Conduit, P.T., and J.W. Raff. 2010. Cnn dynamics drive centrosome size asymmetry to ensure daughter centriole retention in Drosophila neuroblasts. Curr. Biol. 20:2187–2192.

Conduit, P.T., A. Wainman, and J.W. Raff. 2015. Centrosome function and assembly in animal cells. Nat Rev Mol Cell Biol. 16:611–624.

Doe, C.Q. 2008. Neural stem cells: balancing self-renewal with differentiation. Development. 135:1575–1587.

Doe, C.Q., Q. Chu-LaGraff, D.M. Wright, and M.P. Scott. 1991. The prospero gene specifies cell fates in the Drosophila central nervous system. Cell. 65:451–464.

Eliscovich, C., I. Peset, I. Vernos, and R. Mendez. 2008. Spindle-localized CPE-mediated translation controls meiotic chromosome segregation. Nat Cell Biol. 10:858–865.

Fan, Y., and A. Bergmann. 2010. The cleaved-Caspase-3 antibody is a marker of Caspase-9-like DRONC activity in Drosophila. Cell Death Differ. 17:534–539.

Gallaud, E., A. Ramdas Nair, N. Horsley, A. Monnard, P. Singh, T.T. Pham, D. Salvador Garcia, A. Ferrand, and C. Cabernard. 2020. Dynamic centriolar localization of Polo and Centrobin in early mitosis primes centrosome asymmetry. PLoS Biol. 18:e3000762.

Gambarotto, D., C. Pennetier, J.M. Ryniawec, D.W. Buster, D. Gogendeau, A. Goupil, M. Nano, A. Simon, D. Blanc, V. Racine, Y. Kimata, G.C. Rogers, and R. Basto. 2019. Plk4 Regulates Centriole Asymmetry and Spindle Orientation in Neural Stem Cells. Dev Cell. 50:11–24 e10.

Giet, R., D. McLean, S. Descamps, M.J. Lee, J.W. Raff, C. Prigent, and D.M. Glover. 2002. Drosophila Aurora A kinase is required to localize D-TACC to centrosomes and to regulate astral microtubules. J. Cell Biol. 156:437–451.

Gould, R.R., and G.G. Borisy. 1977. The pericentriolar material in Chinese hamster ovary cells nucleates microtubule formation. J Cell Biol. 73:601–615.

Groisman, I., Y.S. Huang, R. Mendez, Q. Cao, W. Theurkauf, and J.D. Richter. 2000. CPEB, maskin, and cyclin B1 mRNA at the mitotic apparatus: implications for local translational control of cell division. Cell. 103:435–447.

Hafer, N., S. Xu, K.M. Bhat, and P. Schedl. 2011. The Drosophila CPEB protein Orb2 has a novel expression pattern and is important for asymmetric cell division and nervous system function. Genetics. 189:907–921.

Hervas, R., L. Li, A. Majumdar, C. Fernandez-Ramirez Mdel, J.R. Unruh, B.D. Slaughter, A. Galera-Prat, E. Santana, M. Suzuki, Y. Nagai, M. Bruix, S. Casas-Tinto, M. Menendez, D.V. Laurents, K. Si, and M. Carrion-Vazquez. 2016. Molecular Basis of Orb2 Amyloidogenesis and Blockade of Memory Consolidation. PLoS Biol. 14:e1002361.

Huang, Y.S., M.C. Kan, C.L. Lin, and J.D. Richter. 2006. CPEB3 and CPEB4 in neurons: analysis of RNA-binding specificity and translational control of AMPA receptor GluR2 mRNA. EMBO J. 25:4865–4876.

Inaba, M., Z.G. Venkei, and Y.M. Yamashita. 2015. The polarity protein Baz forms a platform for the centrosome orientation during asymmetric stem cell division in the Drosophila male germline. eLife. 4.

Januschke, J., and C. Gonzalez. 2010. The interphase microtubule aster is a determinant of asymmetric division orientation in Drosophila neuroblasts. J. Cell Biol. 188:693–706.

Januschke, J., S. Llamazares, J. Reina, and C. Gonzalez. 2011. Drosophila neuroblasts retain the daughter centrosome. Nat. Commun. 2:243.

Januschke, J., J. Reina, S. Llamazares, T. Bertran, F. Rossi, J. Roig, and C. Gonzalez. 2013. Centrobin controls mother-daughter centriole asymmetry in Drosophila neuroblasts. Nat Cell Biol.

Keleman, K., S. Kruttner, M. Alenius, and B.J. Dickson. 2007. Function of the Drosophila CPEB protein Orb2 in long-term courtship memory. Nat Neurosci. 10:1587–1593.

Khan, M.R., L. Li, C. Perez-Sanchez, A. Saraf, L. Florens, B.D. Slaughter, J.R. Unruh, and K. Si. 2015. Amyloidogenic Oligomerization Transforms Drosophila Orb2 from a Translation Repressor to an Activator. Cell. 163:1468–1483.

Khodjakov, A., and C.L. Rieder. 1999. The sudden recruitment of gamma-tubulin to the centrosome at the onset of mitosis and its dynamic exchange throughout the cell cycle, do not require microtubules. J. Cell Biol. 146:585–596.

Knoblich, J.A., L.Y. Jan, and Y.N. Jan. 1995. Asymmetric segregation of Numb and Prospero during cell division. Nature. 377:624–627.

Kraut, R., W. Chia, L.Y. Jan, Y.N. Jan, and J.A. Knoblich. 1996. Role of inscuteable in orienting asymmetric cell divisions in Drosophila. Nature. 383:50–55.

Kruttner, S., B. Stepien, J.N. Noordermeer, M.A. Mommaas, K. Mechtler, B.J. Dickson, and K. Keleman. 2012. Drosophila CPEB Orb2A mediates memory independent of Its RNA-binding domain. Neuron. 76:383–395.

Kuchinke, U., F. Grawe, and E. Knust. 1998. Control of spindle orientation in Drosophila by the Par-3-related PDZ-domain protein Bazooka. Curr Biol. 8:1357–1365.

Lai, S.L., and C.Q. Doe. 2014. Transient nuclear Prospero induces neural progenitor quiescence. eLife. 3.

Larkin, A., S.J. Marygold, G. Antonazzo, H. Attrill, G. Dos Santos, P.V. Garapati, J.L. Goodman, L.S. Gramates, G. Millburn, V.B. Strelets, C.J. Tabone, J. Thurmond, and C. FlyBase. 2021. FlyBase: updates to the Drosophila melanogaster knowledge base. Nucleic Acids Res. 49:D899–D907.

Lee, M.J., F. Gergely, K. Jeffers, S.Y. Peak-Chew, and J.W. Raff. 2001. Msps/XMAP215 interacts with the centrosomal protein D-TACC to regulate microtubule behaviour. Nat Cell Biol. 3:643–649.

Lerit, D.A., K.M. Plevock, and N.M. Rusan. 2014. Live imaging of Drosophila larval neuroblasts. J Vis Exp.

Lerit, D.A., and N.M. Rusan. 2013. PLP inhibits the activity of interphase centrosomes to ensure their proper segregation in stem cells. J. Cell Biol. 202:1013–1022.

Licatalosi, D.D., A. Mele, J.J. Fak, J. Ule, M. Kayikci, S.W. Chi, T.A. Clark, A.C. Schweitzer, J.E. Blume, X. Wang, J.C. Darnell, and R.B. Darnell. 2008. HITS-CLIP yields genome-wide insights into brain alternative RNA processing. Nature. 456:464–469.

Link, N., H. Chung, A. Jolly, M. Withers, B. Tepe, B.R. Arenkiel, P.S. Shah, N.J. Krogan, H. Aydin, B.B. Geckinli, T. Tos, S. Isikay, B. Tuysuz, G.H. Mochida, A.X. Thomas, R.D. Clark, G.M. Mirzaa, J.R. Lupski, and H.J. Bellen. 2019. Mutations in ANKLE2, a ZIKA Virus Target, Disrupt an Asymmetric Cell Division Pathway in Drosophila Neuroblasts to Cause Microcephaly. Dev Cell. 51:713–729 e716.

Martinez-Campos, M., R. Basto, J. Baker, M. Kernan, and J.W. Raff. 2004. The Drosophila pericentrin-like protein is essential for cilia/flagella function, but appears to be dispensable for mitosis. J. Cell Biol. 165:673–683.

Mastushita-Sakai, T., E. White-Grindley, J. Samuelson, C. Seidel, and K. Si. 2010. Drosophila Orb2 targets genes involved in neuronal growth, synapse formation, and protein turnover. Proc Natl Acad Sci U S A. 107:11987–11992.

Megraw, T.L., K. Li, L.R. Kao, and T.C. Kaufman. 1999. The centrosomin protein is required for centrosome assembly and function during cleavage in Drosophila. Development. 126:2829–2839.

Megraw, T.L., J.T. Sharkey, and R.S. Nowakowski. 2011. Cdk5rap2 exposes the centrosomal root of microcephaly syndromes. Trends Cell Biol. 21:470–480.

Mi, H., D. Ebert, A. Muruganujan, C. Mills, L.P. Albou, T. Mushayamaha, and P.D. Thomas. 2021. PANTHER version 16: a revised family classification, tree-based classification tool, enhancer regions and extensive API. Nucleic Acids Res. 49:D394–D403.

Pascual, R., C. Segura-Morales, M. Omerzu, N. Bellora, E. Belloc, C.L. Castellazzi, O. Reina, E. Eyras, M.M. Maurice, A. Millanes-Romero, and R. Mendez. 2020. mRNA spindle localization and mitotic translational regulation by CPEB1 and CPEB4. RNA. 27:291–302.

Ramdas Nair, A., P. Singh, D. Salvador Garcia, D. Rodriguez-Crespo, B. Egger, and C. Cabernard. 2016. The Microcephaly-Associated Protein Wdr62/CG7337 Is Required to Maintain Centrosome Asymmetry in Drosophila Neuroblasts. Cell Rep. 14:1100–1113.

Rebollo, E., P. Sampaio, J. Januschke, S. Llamazares, H. Varmark, and C. Gonzalez. 2007. Functionally unequal centrosomes drive spindle orientation in asymmetrically dividing Drosophila neural stem cells. Dev. Cell. 12:467–474.

Robinson, B.V., V. Faundez, and D.A. Lerit. 2020. Understanding microcephaly through the study of centrosome regulation in Drosophila neural stem cells. Biochem Soc Trans. 48:2101–2115.

Rusan, N.M., and M. Peifer. 2007. A role for a novel centrosome cycle in asymmetric cell division. J. Cell Biol. 177:13–20.

Schindelin, J., I. Arganda-Carreras, E. Frise, V. Kaynig, M. Longair, T. Pietzsch, S. Preibisch, C. Rueden, S. Saalfeld, B. Schmid, J.Y. Tinevez, D.J. White, V. Hartenstein, K. Eliceiri, P. Tomancak, and A. Cardona. 2012. Fiji: an open-source platform for biological-image analysis. Nature methods. 9:676–682.

Schober, M., M. Schaefer, and J.A. Knoblich. 1999. Bazooka recruits Inscuteable to orient asymmetric cell divisions in Drosophila neuroblasts. Nature. 402:548–551.

Shen, C.P., J.A. Knoblich, Y.M. Chan, M.M. Jiang, L.Y. Jan, and Y.N. Jan. 1998. Miranda as a multidomain adapter linking apically localized Inscuteable and basally localized Staufen and Prospero during asymmetric cell division in Drosophila. Genes Dev. 12:1837–1846.

Siegrist, S.E., and C.Q. Doe. 2005. Microtubule-induced Pins/Galphai cortical polarity in Drosophila neuroblasts. Cell. 123:1323–1335.

Siegrist, S.E., and C.Q. Doe. 2006. Extrinsic cues orient the cell division axis in Drosophila embryonic neuroblasts. Development. 133:529–536.

Siller, K.H., C. Cabernard, and C.Q. Doe. 2006. The NuMA-related Mud protein binds Pins and regulates spindle orientation in Drosophila neuroblasts. Nat. Cell Biol. 8:594–600.

Singh, P., A. Ramdas Nair, and C. Cabernard. 2014. The centriolar protein Bld10/Cep135 is required to establish centrosome asymmetry in Drosophila neuroblasts. Curr Biol. 24:1548–1555.

Stepien, B.K., C. Oppitz, D. Gerlach, U. Dag, M. Novatchkova, S. Kruttner, A. Stark, and K. Keleman. 2016. RNA-binding profiles of Drosophila CPEB proteins Orb and Orb2. Proc Natl Acad Sci U S A. 113:E7030–E7038.

Sunkel, C.E., and D.M. Glover. 1988. polo, a mitotic mutant of Drosophila displaying abnormal spindle poles. J Cell Sci. 89 (Pt 1):25–38.

Thornton, G.K., and C.G. Woods. 2009. Primary microcephaly: do all roads lead to Rome? Trends in genetics : TIG. 25:501–510.

Truman, J.W., and M. Bate. 1988. Spatial and temporal patterns of neurogenesis in the central nervous system of Drosophila melanogaster. Developmental Biology. 125:145–157.

Vaessin, H., E. Grell, E. Wolff, E. Bier, L.Y. Jan, and Y.N. Jan. 1991. prospero is expressed in neuronal precursors and encodes a nuclear protein that is involved in the control of axonal outgrowth in Drosophila. Cell. 67:941–953.

Vaizel-Ohayon, D., and E.D. Schejter. 1999. Mutations in centrosomin reveal requirements for centrosomal function during early Drosophila embryogenesis. Curr. Biol. 9:889–898.

Varmark, H., S. Llamazares, E. Rebollo, B. Lange, J. Reina, H. Schwarz, and C. Gonzalez. 2007. Asterless is a centriolar protein required for centrosome function and embryo development in Drosophila. Curr. Biol. 17:1735–1745.

Wang, C., S. Li, J. Januschke, F. Rossi, Y. Izumi, G. Garcia-Alvarez, S.S. Gwee, S.B. Soon, H.K. Sidhu, F. Yu, F. Matsuzaki, C. Gonzalez, and H. Wang. 2011. An ana2/ctp/mud complex regulates spindle orientation in Drosophila neuroblasts. Dev Cell. 21:520–533.

Wodarz, A., A. Ramrath, U. Kuchinke, and E. Knust. 1999. Bazooka provides an apical cue for Inscuteable localization in Drosophila neuroblasts. Nature. 402:544–547.

Xu, S., N. Hafer, B. Agunwamba, and P. Schedl. 2012. The CPEB protein Orb2 has multiple functions during spermatogenesis in Drosophila melanogaster. PLoS genetics. 8:e1003079.

Xu, S., S. Tyagi, and P. Schedl. 2014. Spermatid cyst polarization in Drosophila depends upon apkc and the CPEB family translational regulator orb2. PLoS genetics. 10:e1004380.

Yamashita, Y.M., D.L. Jones, and M.T. Fuller. 2003. Orientation of asymmetric stem cell division by the APC tumor suppressor and centrosome. Science. 301:1547–1550.

