## Supplementary figures and images for "RNA-binding protein Orb2 causes microcephaly and supports centrosome asymmetry in *Drosophila* neural stem cells"

### Supplemental Figure 1

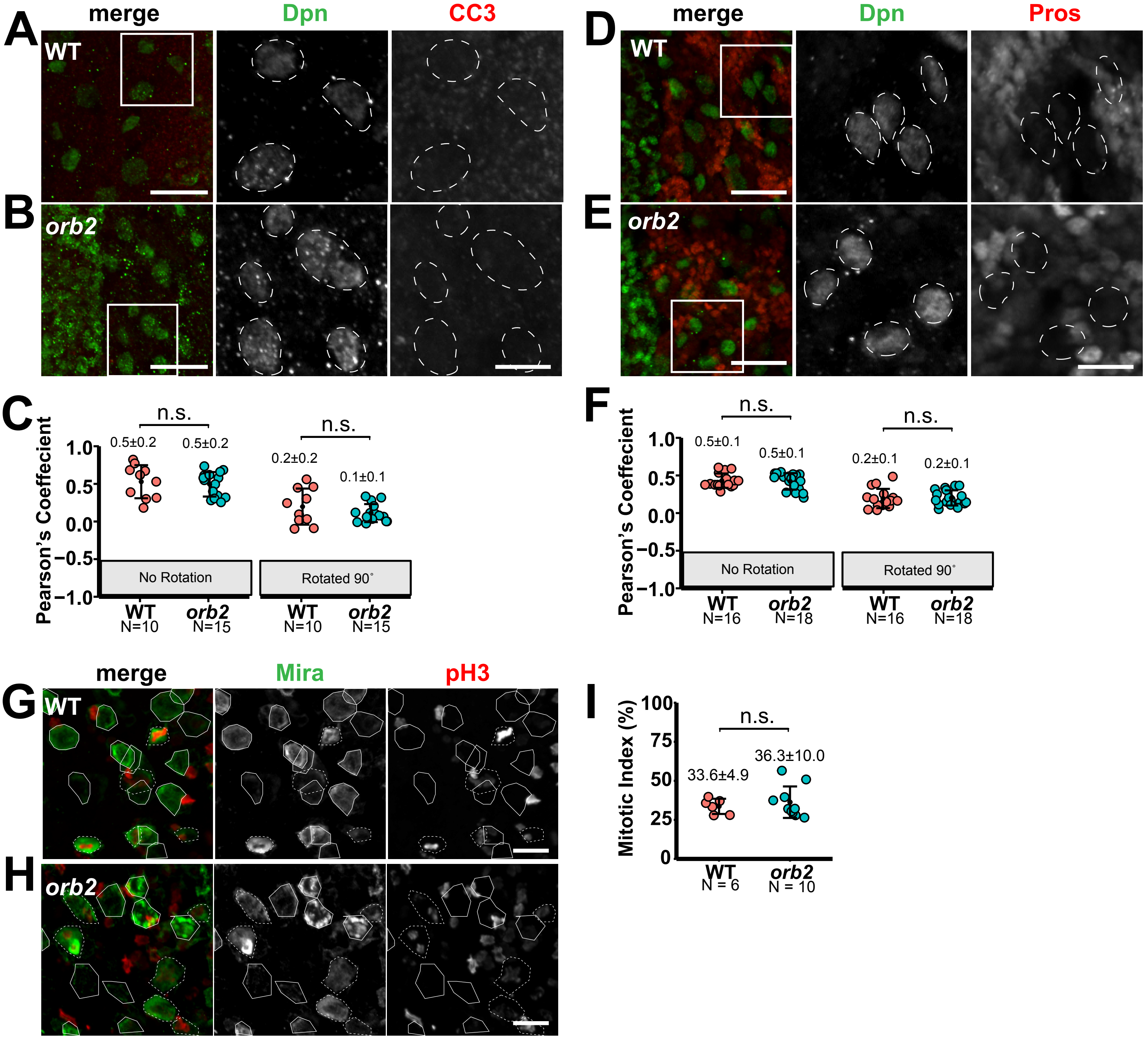


Robinson et al.

Figure S1. NSC differentiation and survival in WT vs. *orb2* mutants

### Supplemental Figure 2

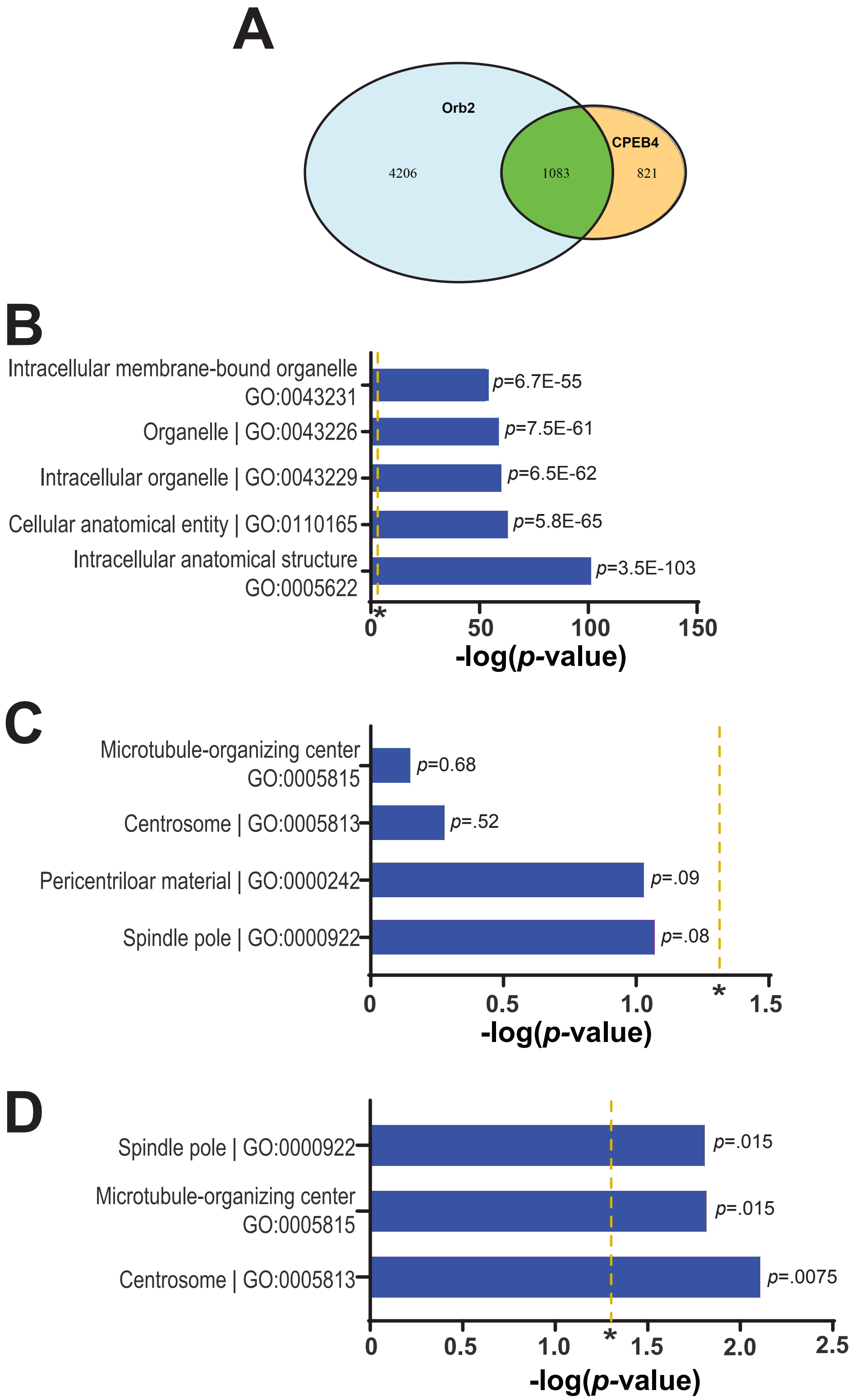


Robinson et al.

Figure S2. Ontological analysis of Orb2 and human CPEB4
